## Supplementary Figures for "*Aeromonas hydrophila* CobQ is a new type of NAD^+^- and Zn^2+^-independent protein lysine deacetylase"

**Figure S12. Characteristics of the overexpressed and purified recombinant target proteins.** The PCR products (right) and SDS-PAGE results of the target recombinant or site-directed acetylated proteins. (A) ArcA-2 and ArcA-2-K26; (B) SUN, SUN-K103, and SUN-K148; (C) ENO and ENO-K195.

**Figure S1**

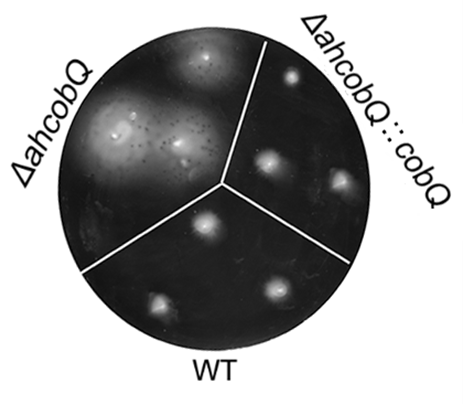

**Figure S2**

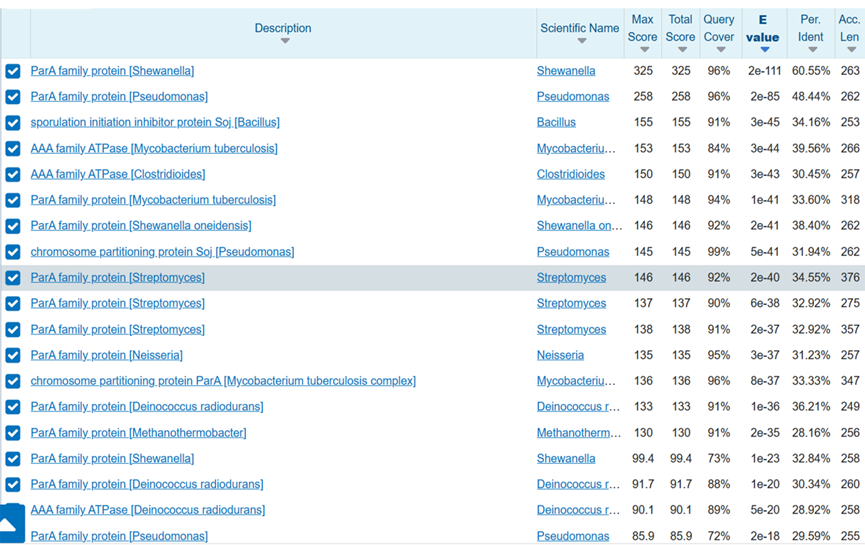

**Figure S3**

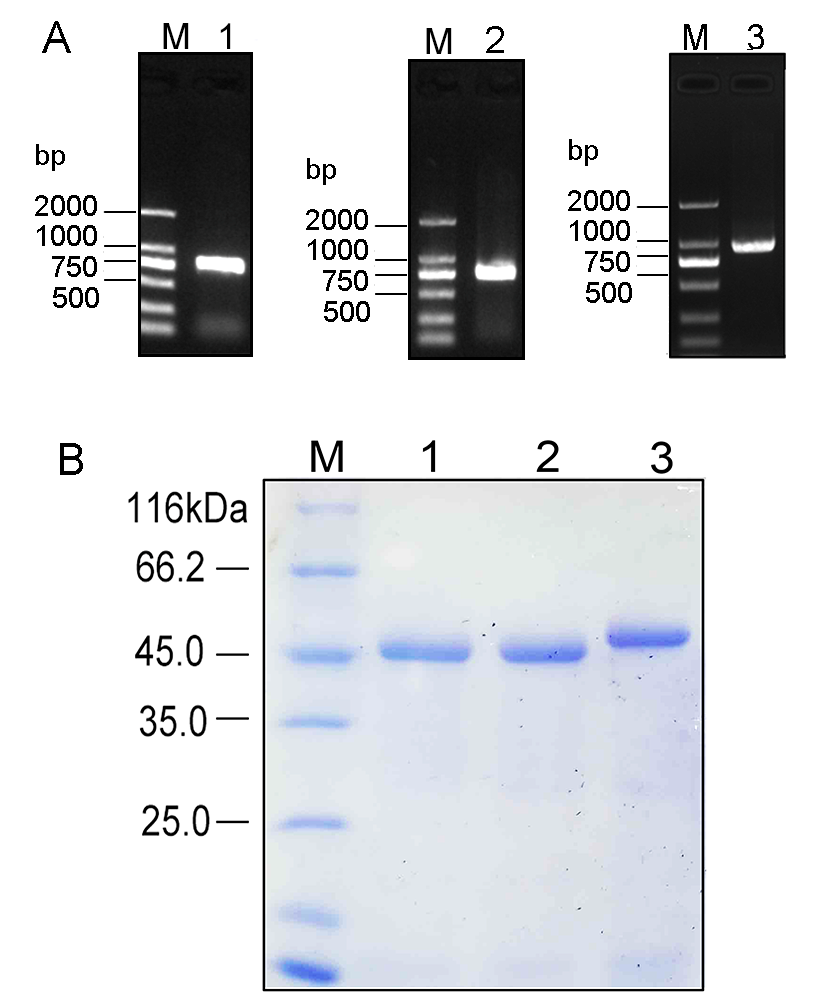

**Figure S4**

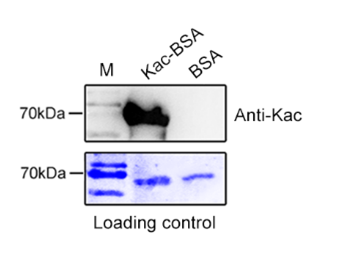

**Figure S**
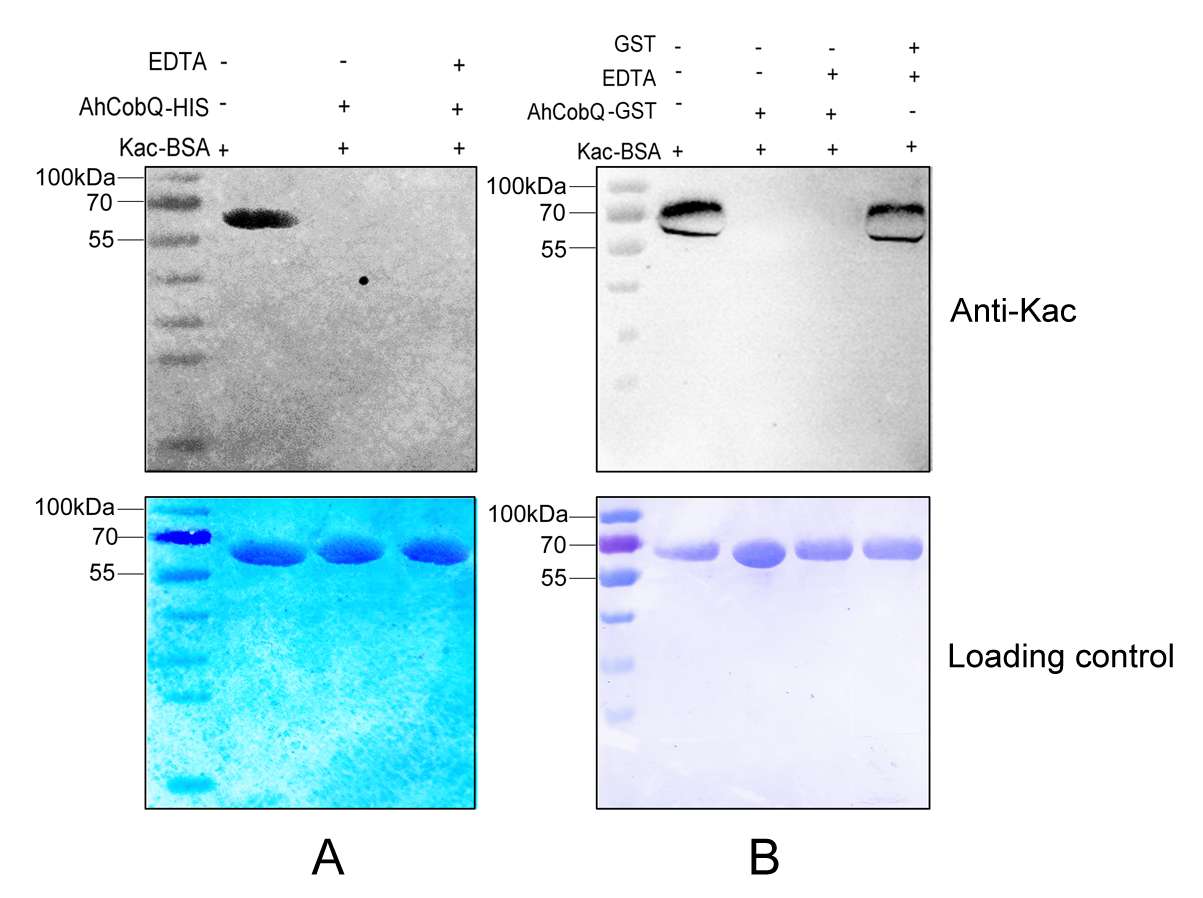
**5**

**Figure S6**

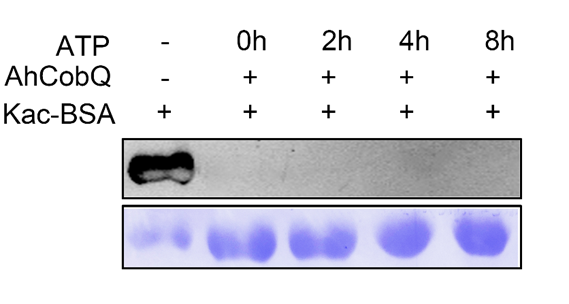

**Figure S7**

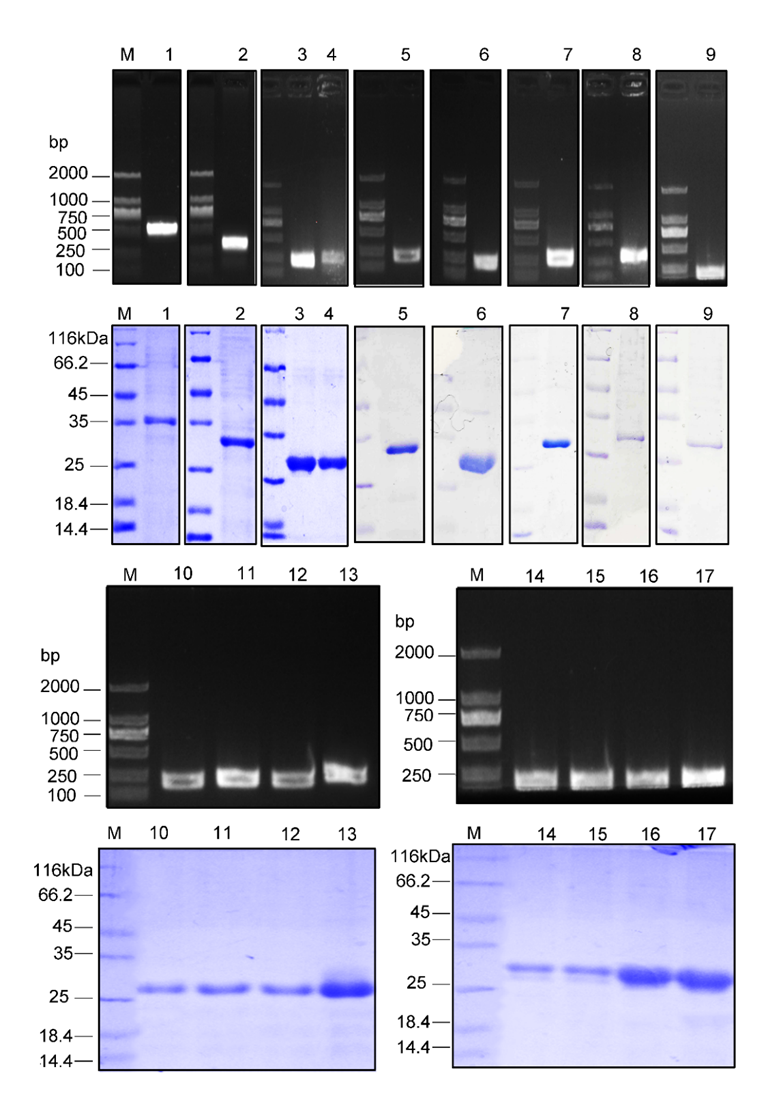

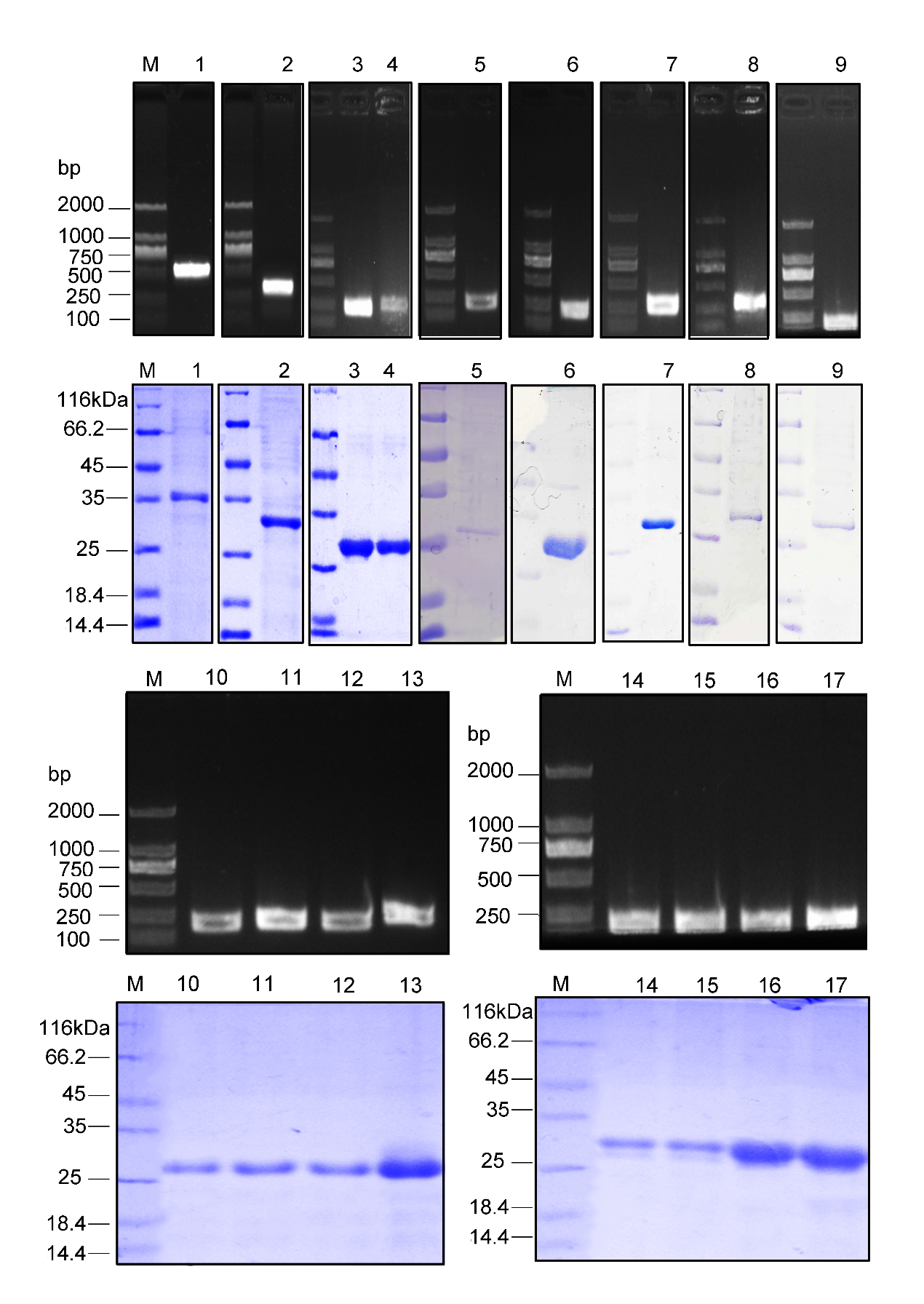

**Figure S8**

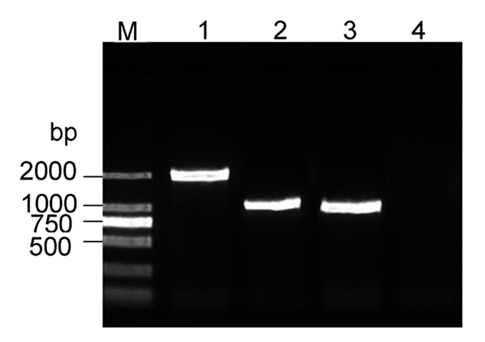

**Figure S9**

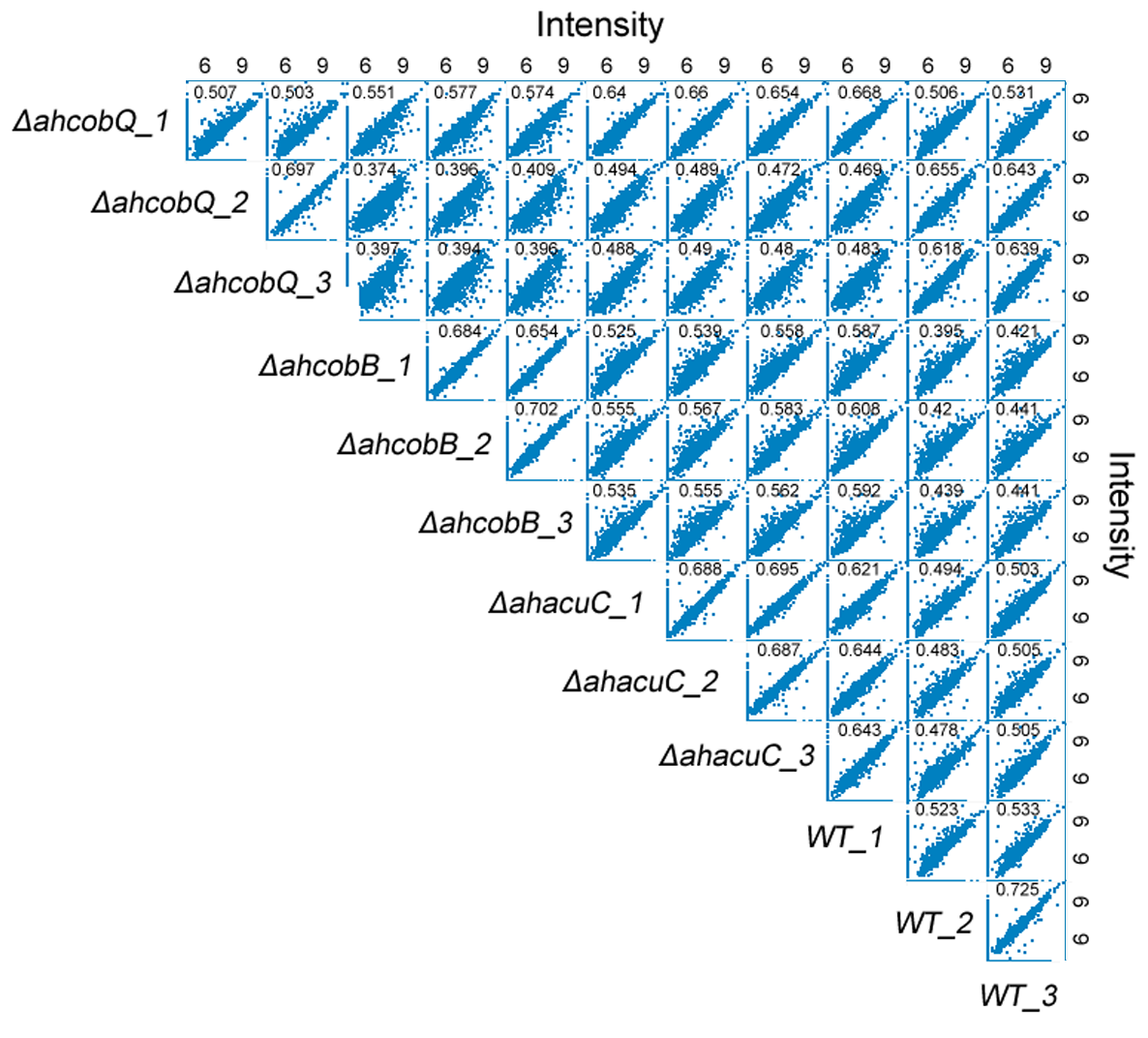

**Figure S10**

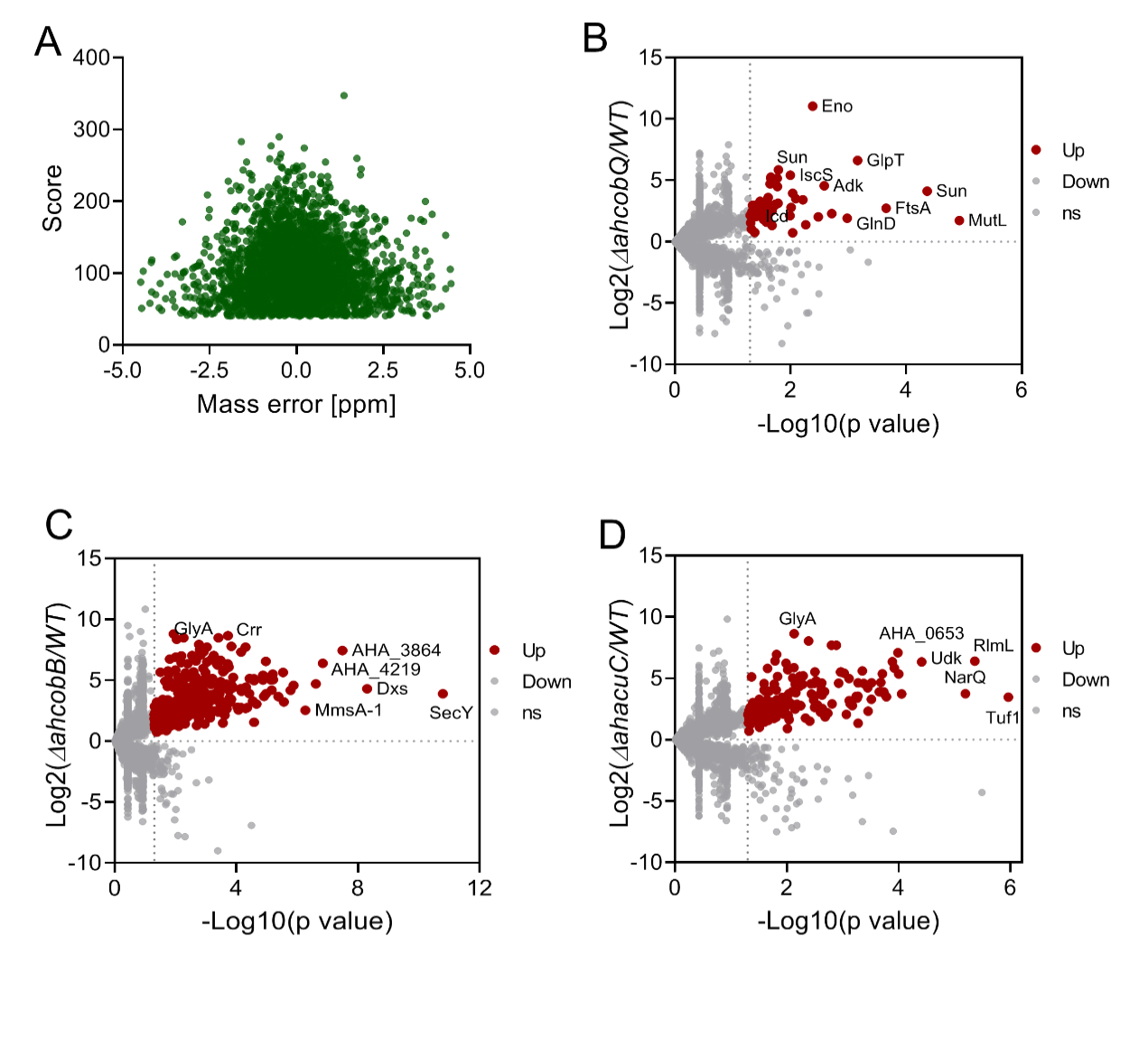

**Figure S11**

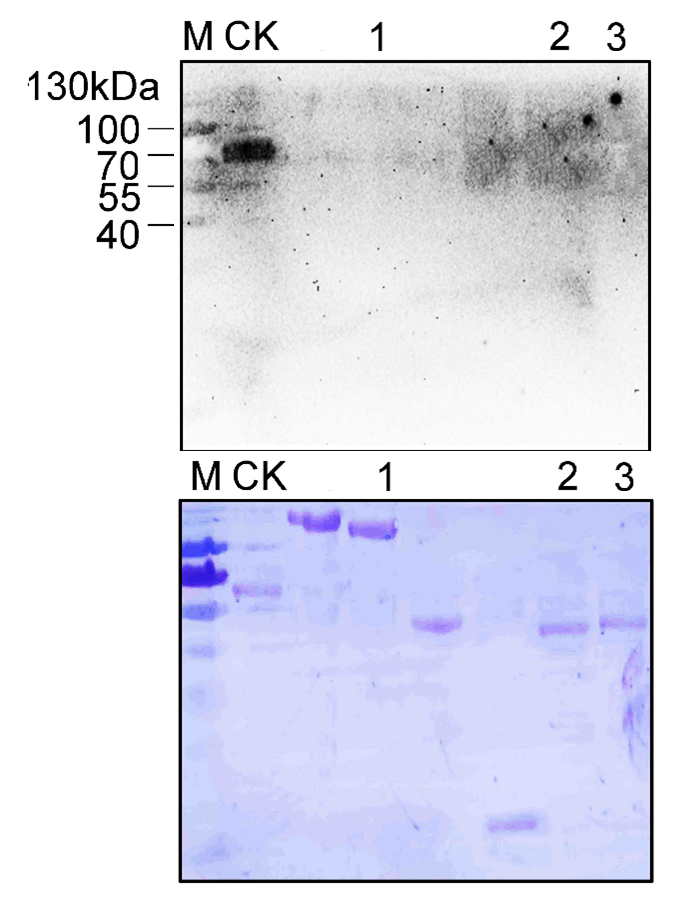

**Figure S12**

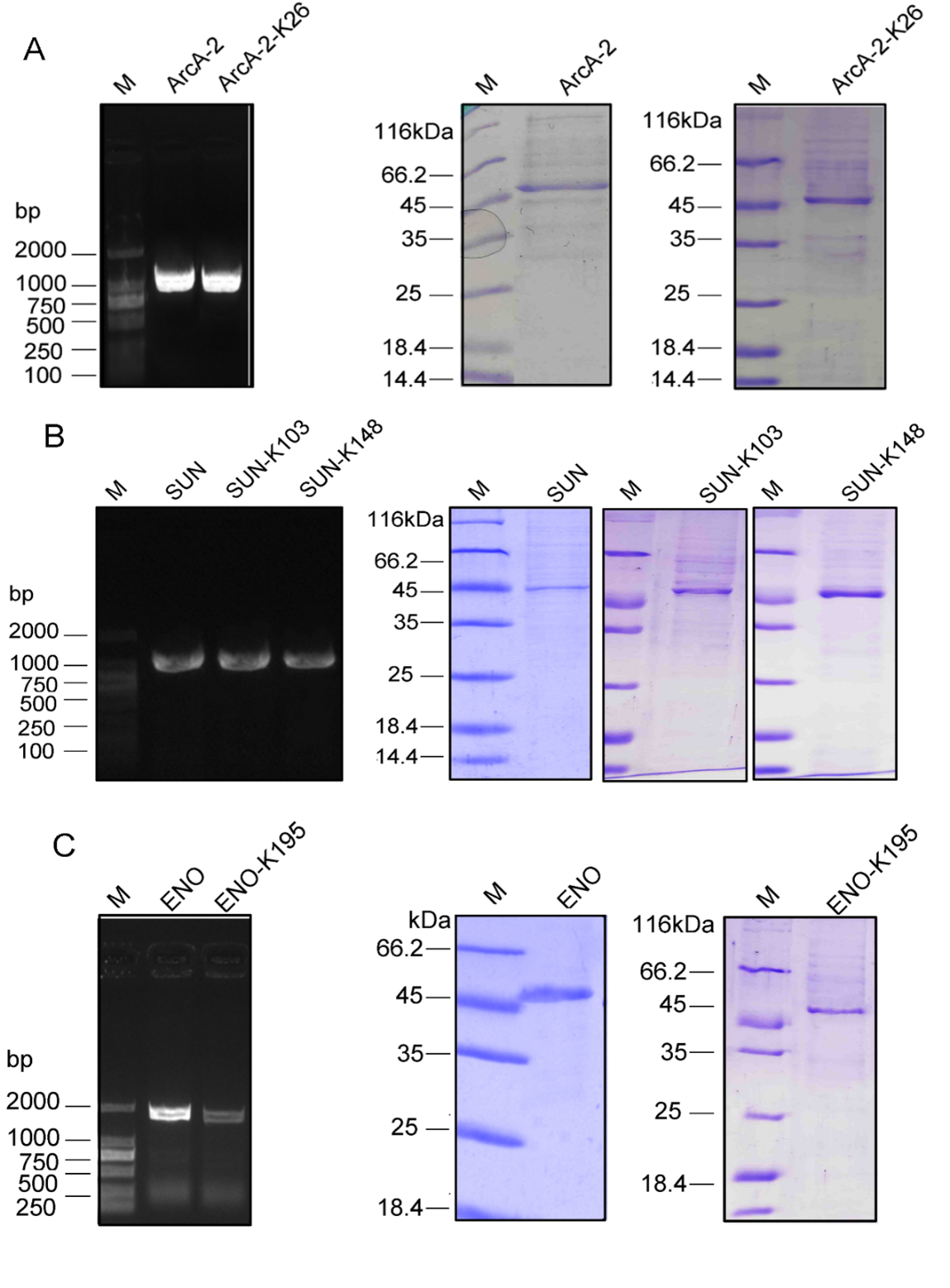
