## Supplementary Tables for "*Aeromonas hydrophila* CobQ is a new type of NAD^+^- and Zn^2+^-independent protein lysine deacetylase"

**Table S1. Selected Kac peptide quantification among Kac-BSA and Kac-BSA incubated with CobQ and BSA without acetylation, by LC MS/MS.**

**Table S2. Acetylation-modified upregulated proteins of AhCobQ-deleted strains and the identification of residue positions by LC MS/MS.**

**Table S3. Chemicals and reagents used in this study.**

**Table S4. Critical commercial assay kits used in this study.**

**Table S5. Bacterial strains and plasmids used in this study.**

**Table S6. Primer pairs used in this study.**

**Table S1. Selected Kac peptide** **quantification of Kac-BSA, Kac-BSA incubated with CobQ, and BSA without acetylation by LC MS/MS**

| Kac peptide in BSA | Score | Intensity  Kac-BSA+CobQ | Intensity Kac-BSA | Intensity BSA |
| --- | --- | --- | --- | --- |
| LVNELTEFAK*TCVADESHAGCEK | 329.06 | 0 | 5.68E+07 | 0 |
| YNGVFQECCQAEDK*GACLLPK | 266.15 | 0 | 2.93E+08 | 0 |
| NYQEAK*DAFLGSFLYEYSR | 208.92 | 0 | 2.63E+08 | 0 |
| EYEATLEECCAK*DDPHACYSTVFDK | 194.62 | 0 | 1.92E+08 | 0 |
| LKPDPNTLCDEFK*ADEK | 182.52 | 0 | 2.52E+08 | 0 |
| ECCDK*PLLEK | 151.98 | 0 | 3.61E+08 | 0 |
| FWGK*YLYEIAR | 141.08 | 0 | 2.13E+08 | 0 |
| RHPYFYAPELLYYANK*YNGVFQECCQAEDK | 131.5 | 0 | 9.07E+07 | 0 |
| DAIPENLPPLTADFAEDK*DVCK | 127.75 | 0 | 3.62E+08 | 0 |
| ATEEQLK*TVMENFVAFVDK | 126.63 | 0 | 8.33E+08 | 0 |
| DVCK*NYQEAK | 124.82 | 0 | 3.38E+08 | 0 |
| K*FWGKYLYEIAR | 119.45 | 0 | 41218000 | 0 |
| AEFVEVTK*LVTDLTK | 101.53 | 0 | 1.28E+09 | 0 |
| CCAADDKEACFAVEGPK*LVVSTQTALA | 62.1 | 0 | 1.15E+08 | 0 |
| QEPERNECFLSHK*DDSPDLPK | 46.027 | 0 | 9.40E+07 | 0 |

Note: asterisk indicates Kac; best Maxquant Kac peptide score > 35 and localization probability > 0.75; BSA: bovine serum albumin.

**Table S2. Acetylation-modified upregulated proteins of AhCobQ-deleted strains and the identification of residue positions by LC MS/MS**

| Protein ID | Gene ID | Description | Positions (K) | Ratio (log2) | P-value |
| --- | --- | --- | --- | --- | --- |
| A0KGH3 | *eno* | Enolase | 195 | 11.04 | 0.004 |
| A0KIT5 | *glpT* | Glycerol-3-phosphate transporter | 152 | 6.62 | 0.001 |
| A0KEW8 | *sun* | Ribosomal RNA small subunit methyltransferase B | 148 | 5.85 | 0.016 |
| A0KJ32 | *iscS* | Cysteine desulfurase IscS | 93 | 5.41 | 0.010 |
| A0KMX2 | *tkt* | Transketolase | 302 | 5.26 | 0.022 |
| A0KPG3 | *recA* | Protein RecA | 279 | 5.18 | 0.022 |
| A0KG67 | *rpsF* | 30S ribosomal protein S6 | 104 | 5.17 | 0.017 |
| A0KF24 | *rplB* | 50S ribosomal protein L2 | 18 | 4.71 | 0.023 |
| A0KL52 | *adk* | Adenylate kinase | 48 | 4.56 | 0.003 |
| A0KMP9 | *rdgC* | Recombination-associated protein RdgC | 46 | 4.50 | 0.017 |
| A0KEW8 | *sun* | Ribosomal RNA small subunit methyltransferase B | 103 | 4.11 | 0.000 |
| A0KKE0 | *hrpA* | ATP-dependent helicase HrpA | 816 | 3.95 | 0.009 |
| A0KP36 | *cysD* | Sulfate adenylyltransferase subunit 2 | 115 | 3.60 | 0.024 |
| A0KHG1 | *glnD* | Bifunctional uridylyltransferase/uridylyl-removing enzyme | 461 | 3.51 | 0.008 |
| A0KHF8 | *ptsG* | PTS system, glucose-specific IIBC component | 380 | 3.41 | 0.006 |
| A0KKQ5 | *nrdA* | Ribonucleoside-diphosphate reductase | 718 | 3.30 | 0.034 |
| A0KKE0 | *hrpA* | ATP-dependent helicase HrpA | 1089 | 3.13 | 0.016 |
| A0KF44 | *rpsD* | 30S ribosomal protein S4 | 156 | 3.12 | 0.024 |
| A0KPV1 | *AHA_3863* | Pyruvate dehydrogenase E1 component | 507 | 3.06 | 0.018 |
| A0KNE8 | *glmM* | Phosphoglucosamine mutase | 34 | 2.98 | 0.041 |
| A0KEE9 | *gyrB* | DNA gyrase subunit B | 331 | 2.97 | 0.019 |
| A0KP36 | *cysD* | Sulfate adenylyltransferase subunit 2 | 82 | 2.96 | 0.045 |
| A0KLH8 | *AHA_2620* | CstA protein | 29 | 2.80 | 0.035 |
| A0KQH8 | *galK* | Galactokinase | 101 | 2.78 | 0.010 |
| A0KPW9 | *ftsA* | Cell division protein FtsA | 320 | 2.72 | 0.000 |
| A0KGI6 | *dnaG* | DNA primase | 241 | 2.70 | 0.029 |
| A0KQA4 | *rpoC* | DNA-directed RNA polymerase subunit beta | 74 | 2.57 | 0.041 |
| A0KJ75 | *AHA_1791* | RNA-binding protein | 81 | 2.52 | 0.021 |
| A0KKJ6 | *AHA_2274* | PhaF | 75 | 2.51 | 0.045 |
| A0KLU0 | *AHA_2733* | Phospho-2-dehydro-3-deoxyheptonate aldolase | 271 | 2.50 | 0.034 |
| A0KKH6 | *AHA_2253* | 3-Oxoacyl-[acyl-carrier-protein] synthase 2 | 329 | 2.49 | 0.025 |
| A0KM13 | *AHA_2809* | Putative transporter | 403 | 2.47 | 0.046 |
| A0KEJ9 | *glyS* | Glycine--tRNA ligase beta subunit | 112 | 2.47 | 0.029 |
| A0KK84 | *AHA_2161* | Aspartate/ornithine carbamoyltransferase, Asp/Orn binding domain family | 307 | 2.36 | 0.023 |
| A0KEL7 | *AHA_0149* | Acyl-CoA synthetase | 797 | 2.29 | 0.002 |
| A0KIV6 | *AHA_1671* | Polyphosphate kinase 2 | 172 | 2.27 | 0.034 |
| A0KJR9 | *guaB* | Inosine-5'-monophosphate dehydrogenase | 110 | 2.15 | 0.050 |
| A0KI76 | *dsdA* | Probable D-serine dehydratase | 118 | 2.13 | 0.044 |
| A0KN87 | *leuS* | Leucine--tRNA ligase | 561 | 2.12 | 0.010 |
| A0KI74 | *icd* | Isocitrate dehydrogenase [NADP] | 388 | 2.07 | 0.039 |
| A0KND9 | *pnp* | Polyribonucleotide nucleotidyltransferase | 523 | 2.03 | 0.003 |
| A0KHG1 | *glnD* | Bifunctional uridylyltransferase/uridylyl-removing enzyme | 205 | 1.92 | 0.001 |
| A0KN82 | *AHA_3242* | PhoH family protein | 219 | 1.83 | 0.025 |
| A0KN37 | *AHA_3197* | Methyl-accepting chemotaxis protein | 262 | 1.81 | 0.029 |
| A0KGR9 | *mutL* | DNA mismatch repair protein MutL | 200 | 1.73 | 0.000 |
| A0KNL1 | *cysH* | Phosphoadenosine phosphosulfate reductase | 193 | 1.60 | 0.026 |
| A0KHG1 | *glnD* | Bifunctional uridylyltransferase/uridylyl-removing enzyme | 686 | 1.54 | 0.049 |
| A0KQG4 | *argF* | Ornithine carbamoyltransferase | 293 | 1.39 | 0.005 |
| A0KJB6 | *AHA_1833* | Uncharacterized protein | 401 | 1.33 | 0.021 |
| A0KQG6 | *arcA-2* | Arginine deiminase | 26 | 1.02 | 0.047 |
| A0KES5 | *AHA_0213* | S-formylglutathione hydrolase | 17 | 0.76 | 0.042 |
| A0KQS4 | *ilvD* | Dihydroxy-acid dehydratase | 320 | 0.73 | 0.009 |

**Table S3. Chemicals and reagents used in this study.**

| Reagent | Company | Lot number |
| --- | --- | --- |
| Yeast extract | Oxoid | Cat# 4304391-02 |
| Tryptone | Oxoid | Cat# 3153896 |
| Sodium chloride | Sinophem | Cat# 10019318 |
| Agar powder | Solarbio | Cat# A8190 |
| Ligase | Vazyme | Cat# C113 |
| 2 x Rapid Taq Master Mix | Vazyme | Cat# P222 |
| 2 x Phanta Max Master Mix | Vazyme | Cat# P515-03 |
| Ampicillin | Aladdin | Cat# 69-52-3 |
| Chloramphenicol | Solarbio | Cat# A8193 |
| Nicotinamide | Sigma Aldrich |  |
| BSA | Sigma Aldrich |  |
| IPTG | Solarbio |  |
| N-acetyllysine | Sigma Aldrich |  |
| NAD^+^ | Sigma Aldrich |  |
| Isocitric acid | Sigma Aldrich |  |
| NADPH | Sigma Aldrich |  |
| NADP^+^ | Sigma Aldrich |  |
| PEP/2-PGE | Sigma Aldrich |  |
| Acetylated modified antibody | Jingjie PTM | PTM-101 |
| HRP Goat-mouse IgG | CWBIO | HA1006 |
| ATP | Sigma Aldrich |  |
| Ni-NTA agarose beads | Yeasen |  |
| C18 ZipTips | Millipore |  |
| Acetonitrile (ACN) | Sigma Aldrich |  |
| Formic acid (FA) | Sigma Aldrich |  |
| Sheep blood | Solarbio |  |
| Skim milk | Wako |  |
| Crystal Violet | Sigma Aldrich |  |
| Tris (pH 8.0) | Sangon |  |
| Dithiothreitol (DTT) | Sigma Aldrich |  |
| Iodoacetamide (IAA) | Sigma Aldrich |  |
| Trypsin | Promega |  |
| PVDF | Bio-RAD |  |
| hPLG | Hepeng |  |

**Table S4. Critical commercial assays kits used in this study.**

| Kits | Company | Lot number |
| --- | --- | --- |
| Plasmid extraction kit | Magen | Cat# P1001-03C |
| Gel recovery kit | Magen | Cat# D2111-03 |
| DNA extraction kit | Magen | Cat# D3146-02 |
| Mut Express II Fast Mutagenesis kit | [TransGen Biotech](https://www.so.com/link?m=bsNdbBdOFsB2/NKScvb0A/QanmX3LKNtx0r2RmBqaUeQg/o4ldsJyT0pBfdlVO6pVusGZcRCtrcMsTqnF1As/QyhwXJN51f+gVZL2Vxc8ZXUd8cjtG9lCe+eKS274lT+kqIUfSI5lVX4=) | Cat# FM111-01 |
| ClonExpress MultiS One Step Cloning kit | Vazyme | C113 |
| ClonExpress II One Step Cloning kit | Vazyme | C112 |

**Table S5. Bacterial strains and plasmids used in this study**

| Strain name | Strain type | | Other information |
| --- | --- | --- | --- |
| A. *hydrophila* ATCC 7966 | Wild type strain (WT) |  | |
| *A. hydrophila* LP-2 | Virulent strain |  | |
| *E.coli* BL21 (DE3) | Transformed host (Overexpression) |  | |
| *E.coli* DH5α | Transformed host (Site-directed mutagenesis) |  | |
| *E.coli* MC1061 | Transformed host (Knockout) |  | |
| *E.coli* S17-1λpir | Transformed host (Knockout) |  | |
| *ΔahcobQ* | Knockout strain |  | |
| *ΔahcobB* | Knockout strain |  | |
| *ΔahacuC* | Knockout strain |  | |
| AhCobQ-6His tag | Overexpression strain | pET-32a | |
| AhCobQ-GST tag | Overexpression strain | pGEX-KG | |
| AhCobB-32a | Overexpression strain | pET-32a | |
| AhAcuC-32a | Overexpression strain | pET-32a | |
| AhCobQ_1–179_ | Overexpression strain | pET-32a | |
| AhCobQ_179–265_ | Overexpression strain | pET-32a | |
| AhCobQ_189–265_ | Overexpression strain | pET-32a | |
| AhCobQ_189–255_ | Overexpression strain | pET-32a | |
| AhCobQ_189–250_ | Overexpression strain | pET-32a | |
| AhCobQ_189–245_ | Overexpression strain | pET-32a | |
| AhCobQ_189–240_ | Overexpression strain | pET-32a | |
| AhCobQ_195–255_ | Overexpression strain | pET-32a | |
| AhCobQ_200–255_ | Overexpression strain | pET-32a | |
| AhCobQ_179–255_ | Overexpression strain | pET-32a | |
| AhCobQ_179–245_ | Overexpression strain | pET-32a | |
| AhCobQ_179–235_ | Overexpression strain | pET-32a | |
| AhCobQ_179–225_ | Overexpression strain | pET-32a | |
| AhCobQ_199–265_ | Overexpression strain | pET-32a | |
| AhCobQ_209–265_ | Overexpression strain | pET-32a | |
| AhCobQ_219–265_ | Overexpression strain | pET-32a | |
| SUN-K103 | Site-directed acetylation strain | pET-21b | |
| SUN-K148 | Site-directed acetylation strain | pET-21b | |
| ENO-K195 | Site-directed acetylation strain | pET-21b | |
| ArcA-2-K26 | Site-directed acetylation strain | pET-21b | |
| ICD-K388 | Site-directed acetylation strain | pET-21b | |
| pET-32a | Expression vector, HIS-tag, Amp^R^ |  | |
| pET-21b | Expression vector, HIS-tag, Amp^R^ |  | |
| pGEX-KG | Expression vector, GST-tag, Amp^R^ |  | |
| pRE112 | Suicide vector, oriVR6Kγ, Cm^R^, sacB |  | |
| pTECH-MbAcK3RS (IPYE) | Site-directed acetylation co-transformation vector |  | |

**Table S6. Primer pairs used in this study**

| Primer name | Oligonucleotide sequence (5′ → 3′) |
| --- | --- |
| Δ*ahcobB*-P1 | cgatcccaagcttcttctagaCTTGGGATAGGTGGTGAACGG |
| Δ*ahcobB*-P2 | ggaatcagtcGCTATTGTGCGACTGGACAGTTG |
| Δ*ahcobB*-P3 | gcacaatagcGACTGATTCCCTCTGGCTACTTCT |
| Δ*ahcobB*-P4 | catgaattcccgggagagctcGCGAGCAGAGCGTCTACCTG |
| Δ*ahcobB*-P5 | TCAGGGACCGCGCATCTCCATCCAG |
| Δ*ahcobB*-P6 | ATGGTGCAGTCAGCGAAACACATCG |
| Δ*ahcobB*-P7 | GCAACCAACAGCAGTTTGT |
| Δ*ahcobB*-P8 | GCCCTGCCGCTGA |
| Δ*ahcobQ*-P1 | catgaattcccgggagagctcCGAGCTGAGCTCAGGTCTGTTG |
| Δ*ahcobQ*-P2 | ctcTGTATCAATCCTGCCTCGTTGTT |
| Δ*ahcobQ*-P3 | gaggcaggattgatacaGAGATACCACCCAAGTTCGGG |
| Δ*ahcobQ*-P4 | cgatcccaagcttcttctagaCTCGGTCAACGGCACCGC |
| Δ*ahcobQ*-P5 | GTGATTGTTTGGACGGTTGCC |
| Δ*ahcobQ*-P6 | GATGTTACCTCATGAGCACGCTG |
| Δ*ahcobQ*-P7 | TGCAGCCACGCAGCGACGGTTTGCT |
| Δ*ahcobQ*-P8 | GGCGCCATTTGACCTGGTCACGATG |
| Δ*ahacuC*-P1 | catgaattcccgggagagctcTGGTTGTCGCCTTTGAAGGC |
| Δ*ahacuC*-P2 | cgataaacggGAAACGAGTGCTCGCGCG |
| Δ*ahacuC*-P3 | cactcgtttcCCGTTTATCGCTTCGTCAATC |
| Δ*ahacuC*-P4 | cgatcccaagcttcttctagaCTGTTTGTGCTGGGCGCC |
| Δ*ahacuC*-P5 | GTGCGACAAGGGGAGCCTG |
| Δ*ahacuC*-P6 | CTAGCCGTAGCGCTTTTTCGCC |
| Δ*ahacuC*-P7 | ACCAACCCCTACGTCGTACAGC |
| Δ*ahacuC*-P8 | GTTCGGCCGCTTACCGGTG |
| AhCobQ-32a-F | gctgatatcggatccgaattcGTGATTGTTTGGACGGTTGCC |
| AhCobQ-32a-R | ctcgagtgcggccgcaagcttTCATGATGTTACCTCATGAGCACG |
| AhCobB-32a-F | gctgatatcggatccgaattcATGGTGCAGTCAGCGAAACAC |
| AhCobB-32a-R | ctcgagtgcggccgcaagcttTCAGGGACCGCGCATCTC |
| AhAcuC-32a-F | gctgatatcggatccgaattcGTGCGACAAGGGGAGCCT |
| AhAcuC-32a-R | ctcgagtgcggccgcaagcttCTAGCCGTAGCGCTTTTTCG |
| AhCobQ_1–179_-F | gctgatatcggatccgaattcGTGATTGTTTGGACGGTTGCC |
| AhCobQ_1–179_-R | ctcgagtgcggccgcaagcttTCAGAACTTCTCCCGCTTGG |
| AhCobQ_179–265_-F | gctgatatcggatccgaattcATGCGTTATACCGTCATTCCCA |
| AhCobQ_179–265_-R | ctcgagtgcggccgcaagcttTCATGATGTTACCTCATGAGCACG |
| AhCobQ_189–265_-F | gctgatatcggatccgaattcATGGACAAGCGAACCCGTGCC |
| AhCobQ_189–265_-R | ctcgagtgcggccgcaagcttTCATGATGTTACCTCATGAGCACG |
| AhCobQ_189–255_-F | gctgatatcggatccgaattcATGGACAAGCGAACCCGTGCC |
| AhCobQ_189–255_-R | ctcgagtgcggccgcaagcttTCACTCTTGCGCATCAAGATAGTTGA |
| AhCobQ_189–250_-F | gctgatatcggatccgaattcATGGACAAGCGAACCCGTGCC |
| AhCobQ_189–250_-R | ctcgagtgcggccgcaagcttTCAATAGTTGAGCAGGGTCTCATAGGC |
| AhCobQ_189–245_-F | gctgatatcggatccgaattcATGGACAAGCGAACCCGTGCC |
| AhCobQ_189–245_-R | ctcgagtgcggccgcaagcttTCACTCATAGGCATAGGTGCCACG |
| AhCobQ_189–240_-F | gctgatatcggatccgaattcATGGACAAGCGAACCCGTGCC |
| AhCobQ_189–240_-R | ctcgagtgcggccgcaagcttTCAGCCACGGCTGCTCGGCGA |
| AhCobQ_195–255_-F | gctgatatcggatccgaattcATGTCGCTGATGACTCTGCAGTCC |
| AhCobQ_195–255_-R | ctcgagtgcggccgcaagcttTCACTCTTGCGCATCAAGATAGTTGA |
| AhCobQ_200–255_-F | gctgatatcggatccgaattcATGCAGTCCATCAAGGAGCAGCAC |
| AhCobQ_200–255_-R | ctcgagtgcggccgcaagcttTCACTCTTGCGCATCAAGATAGTTGA |
| SUN-K103-F | gagaccgtcaacgccgtcTAGctgctcaagggcacctccttgcgc |
| SUN-K103-R | CTAgacggcgttgacggtctcggccaccgcggcgtgagccggaat |
| SUN-K148-F | ccggagtggctcaccTAGcggctgcgccaggcctatccggatgag |
| SUN-K148-R | CTAggtgagccactccgggtggccgagacgaatgctggggacacggt |
| ENO-K195-F | gtgttccacaacctggccTAGgtgctgaagtccaagggctacaac |
| ENO-K195-R | CTAggccaggttgtggaacacttcagcgcccatgcggacagcttctttc |
| ArcA-2-K26-F | cccaacctcagtctgTAGcgtctgactccttccaactgccaggat |
| ArcA-2-K26-R | CTAcagactgaggttggggcggtgcaacatgacacggcgcaatt |
| ICD-K388-F | gcggcgatccgcaacTAGaccgtcacttatgacttcgagcgtctg |
| ICD-K388-R | CTAgttgcggatcgccgcttccatccccttgatgatgagatcggc |
| SUN-F | gctgatatcggatccgaattcATGAAAACACGCGCACAGG |
| SUN-R | ctcgagtgcggccgcaagcttTTACCGCTTGATCAGCTTGGC |
| ENO-F | gctgatatcggatccgaattcATGTCCAAGATCGTTAAAGTGATCG |
| ENO-R | ctcgagtgcggccgcaagcttTTAAGCCTGGTTCTTCACTTCTTTC |
| ArcA-2-F | aatgggtcgggatccgaattcATGAGCAAATTTTATGTAGGTTCTGAA |
| ArcA-2-R | ctcgagtgcggccgcaagcttTTAGATGCCGTCGCGTTCC |
| ICD-F | gctgatatcggatccgaattcATGGAAAGCAAAGTAGTTATCCCG |
| ICD-R | ctcgagtgcggccgcaagcttTTACATCTGGTCGACCATGTCCT |
